# ScatTR: Estimating the Size of Long Tandem Repeat Expansions from Short-Reads

**DOI:** 10.1101/2025.02.15.638440

**Authors:** Rashid Al-Abri, Gamze Gürsoy

## Abstract

Tandem repeats (TRs) are sequences of DNA where two or more base pairs are repeated back-to-back at specific locations in the genome. The expansions of TRs are implicated in over 50 conditions, including Friedreich’s ataxia, autism, and cancer. However, accurately measuring the copy number of TRs is challenging, especially when their expansions are larger than the fragment sizes used in standard short-read genome sequencing. Here we introduce ScatTR, a novel computational method that leverages a maximum likelihood framework to estimate the copy number of large TR expansions from short-read sequencing data. ScatTR calculates the likelihood of different alignments between sequencing reads and reference sequences that represent various TR lengths and employs a Monte Carlo technique to find the best match. In simulated data, ScatTR outperforms state-of-the-art methods, particularly for TRs with longer motifs and those with lengths that greatly exceed typical sequencing fragment sizes. When applied to data from the 1000 Genomes Project, ScatTR detected potential large TR expansions that other methods missed, highlighting its ability to better identify genome-wide characterization of TR variation. ScatTR can be accessed via: https://github.com/g2lab/scattr.

## 1. Introduction

Tandem repeats (TRs) are consecutively repeated nucleotide sequence motifs that are widespread throughout the human genome. Studying TR variation in the human population is important due to their high mutation rates and polymorphic nature, which impact various biological processes such as gene expression and diseases such as neurodegeneration, cancer, neurodevelopmental and psychiatric disorders (Gymrek, Willems, et al. 2016; Zhou et al. 2022; Erwin et al. 2023; Trost et al. 2020; Song et al. 2018). Pathogenic repeat expansions vary widely in size, with size often having a correlation with earlier onset and more severe disease phenotype. For example, a CGG expansion of more than 200 units located in the 5’ untranslated region of the *FMR1* gene is known to cause Fragile-X-associated tremor/ataxia syndrome (FXTAS) (Hunter et al. 1993). Interestingly, the number of repeats correlates with earlier onset of tremor and ataxia (Tassone et al. 2007; Leehey et al. 2008). Expansions in *RFC1*, which cause cerebellar ataxia, neuropathy, and vestibular areflexia syndrome (CANVAS), typically exceed 250 repeats, with larger expansions also linked to earlier onset and more severe symptoms (Currò et al. 2024; Cortese et al. 2019). In contrast, some repeat expansions can be much larger. In Huntington’s disease, CAG expansions in *HTT* can grow somatically from 40 to over 500 repeats, driving neurodegeneration once a critical threshold is surpassed (Caron et al. 1993). Myotonic dystrophy type 1 (DM1), caused by *DMPK* expansions, can range from 50 to more than 3000 units, with disease severity increasing alongside expansion size (Brook et al. 1992). Moreover, an expansion in the intron of *C9orf72* is known to cause amyotrophic lateral sclerosis (ALS) (N Siddique and T Siddique 1993). Its size can exceed 2100 repeat units, with size also correlating with methylation status and age of onset (Gijselinck et al. 2016). Despite their critical role in biological processes and their link to various diseases, the precise lengths of large TR expansions remain underexplored due to significant technical challenges.

Studying repetitive DNA sequences has historically been hindered by the lack of tools. Many genomic studies utilize short read-based whole genome sequencing (WGS), but exclude half of the data with RepeatMasker, a tool that identifies repetitive DNA sequences for removal. Currently, over 56% of the human genome is removed with RepeatMasker (Nishimura 2000). Repeat expansions are extremely difficult to detect due to various reasons (Martorell et al. 1997). They can be highly variable in size and location, making them difficult to detect through simple alignment-based methods. Their sizes can be much larger than a typical fragment length in short-read-based WGS data and usually occur in regions of the genome that are difficult to align. To overcome these challenges, various computational methods have been developed to identify repeat expansions from short-read sequencing data.

Current tools, such as STRling, ExpansionHunter, and GangSTR, use likelihood-based approaches to estimate TR copy number from short-read sequencing data but struggle with accuracy when the TR length exceeds the lengths of the fragments used in short-read sequencing (Dashnow, Pedersen, et al. 2022; Dolzhenko, Deshpande, et al. 2019; Mousavi et al. 2019). Danteintroduces an alternative alignment-free approach that applies Hidden Markov Models (HMMs) to model repeat sequences and corrects for polymerase-induced stutter artifacts. While Dante provides robust genotyping for complex TR loci, its accuracy is similarly limited for TRs that extend beyond the read length (Budiš et al. 2019). However, many pathogenic TRs expand to hundreds of base pairs (Hannan 2018). Moreover, these methods are constrained by the length of the TR unit (*i*.*e*., motif) they can analyze: STRling is restricted to motifs between 2-6 base pairs (bp), whereas ExpansionHunter and GangSTR support repeat unit sizes from 2 to 20 bp. While newer methods like TRGT overcome these limitations by leveraging long-read sequencing data, short-read sequencing data is still the overwhelming majority of currently available sequencing data due to its financial feasibility (Dolzhenko, English, et al. 2023). Thus, there is still a need for a computational method that can accurately estimate TR copy numbers that are larger than fragment lengths.

In this study, we introduce ScatTR, a novel computational method designed to accurately estimate the copy number of large TRs from short-read WGS data. ScatTR utilizes a novel maximum likelihood framework based on the best alignment of reads to reference sequences with variable TR copy numbers (dubbed as “decoy references”) using a combination of golden section search (GSS), Monte Carlo simulations, and simulated annealing. In our benchmarks using simulated WGS data, ScatTR demonstrates superior performance, particularly for TRs with longer repeat motifs and large copy numbers. Furthermore, when applied to real data from the 1000 Genomes Project, ScatTR identifies several large TR expansions that go undetected by existing methods. ScatTR provides a powerful new approach for studying TR variation in health and disease.

## 2 Results

### 2.1 Overview of the ScatTR method

ScatTR estimates the copy number of TR regions in a genome by reformulating copy number estimation as an optimization problem that considers both mapped and unmapped reads from aligned paired-end short-read WGS data. It uses a predefined catalog of TR loci that specifies the chromosome, start and end positions, and the repeat motif for each region of interest. For each TR locus, ScatTR identifies relevant read pairs by scanning the alignment file for reads where at least one mate overlaps the flanking regions or for read pairs—mapped or unmapped—whose sequences show high similarity to the repeat motif. All rotations of the motif are taken into account to ensure sensitivity, since certain repeats can appear differently depending on strand orientation and sequence context. These read pairs, collectively called the “bag of reads,” may include “anchored” in-repeat read (IRR) pairs (in which one mate maps to the flanking region while the other lies within the repeat) and “fully IRR” pairs (in which both mates are inside the repeat). This approach differs from many existing methods by explicitly incorporating fully IRR pairs, which provide valuable information when the repeat region exceeds the sequencing fragment length.

Once the bag of reads is extracted, ScatTR estimates the copy number by defining a “decoy reference” for each candidate TR copy number. A decoy reference is constructed by concatenating the flanking sequences on either side of the repeat with a specified number of repeated motif units. The algorithm then seeks to determine which decoy reference best explains the observed reads. It does so by adopting a maximum likelihood framework: if one treats the repeat copy number as unknown, the most likely value is the one that maximizes the probability of observing the bag of reads (also known as the likelihood). An exact definition of the likelihood of the bag involves summing over the probabilities of every combination of paired read alignments Ghodsi et al. 2013. Because directly summing over all combinations is computationally infeasible, ScatTR approximates the likelihood of the bag by identifying the single “best alignment” of the reads in the bag to the decoy reference for each copy number and using that best alignment to calculate a likelihood value. This best alignment is found via Monte Carlo-based simulated annealing (SA), a process that iteratively proposes new alignments for the reads on the decoy reference and allows occasional acceptance of higher-cost (lower-likelihood) alignments so that the search can escape local minima. The cost function being minimized is the negative log like lihood of the bag, which accounts for three main factors: how well each read’s sequence aligns to the positions it is assigned, how the observed insert sizes compare to the sample’s overall insert size distribution, and how the observed depth compares to the sample’s read depth distribution. The read depth and insert size distributions are derived by sampling random, high-quality regions of the reference-aligned WGS sample to capture the typical patterns one expects to see from a correct alignment of the reads.

After simulating the alignment process and computing the cost for a particular copy number, ScatTR compares the result against other copy numbers to see which one yields the highest likelihood (lowest cost). Note that we also provide a number of optimizations to speed up this process. For example, the scores describing how well reads align to positions that place them fully in the repetitive region are computed once and re-used. Moreover, rather than testing every possible copy number, which would be time-consuming, it narrows the search using golden section search (GSS). GSS assumes the likelihood function is unimodal over a given range, successively shrinking that range in a computationally efficient manner to find its minimum. To define a starting interval for the search, ScatTR calculates a rough closed-form estimate of the copy number based on the sample’s mean read depth and the number of IRRs. Multiple executions of the GSS procedure (together with repeated simulated annealing runs), known as bootstrapping, are then performed for robustness, and a final copy number is chosen based on the aggregated results. This bootstrapping step provides a measure of confidence, allowing ScatTR to report not only the most likely copy number but also a confidence interval Methods for that estimate. Please see for details of the formulations and the algorithm.

### 2.2 ScatTR outperforms existing methods on simulated data

We compared our method against STRling, ExpansionHunter, GangSTR, in addition to a closed-form solution (see 4.3) by calculating the Root Mean Square Error (RMSE) between the predicted and true sum of TR copy numbers for each allele (Dashnow, Pedersen, et al. 2022; Dolzhenko, Bennett, et al. 2020; Mousavi et al. 2019).Specifically, for each true copy number, we compute RMSE across all simulated samples with that same true copy number. A lower RMSE indicates that a method produces more accurate predictions. We sampled a subset of 30 TR loci identified by Tandem Repeats Finder (TRF) with motif lengths between 2-20 bp (Benson 1999) and simulated and reference-aligned short-read paired-end WGS samples with expansions for these loci (Supplemental Table 1). We simulated the samples with 30x coverage and a sequencing error profile based on the Illumina HiSeq X platform (see 4.2). We simulated homozygous and heterozygous expansions in 30 TR loci with copy numbers ranging from 200 to 1000 for a total of 540 samples. We compared the predictions with respect to heterozygosity of TR and repeat motif length. We observe that ScatTR’s predictions yield the lowest RMSE across all tested heterozygous TR copy numbers and 8 out of 9 homozygous TR copy numbers (Figure 1A). Additionally, we observe that the copy number of heterozygous TRs is generally better predicted than those of homozygous TRs in all methods including ScatTR. Moreover, we show that the RMSE of ScatTR’s predictions is consistent across different TR copy numbers.

**Figure 1:**
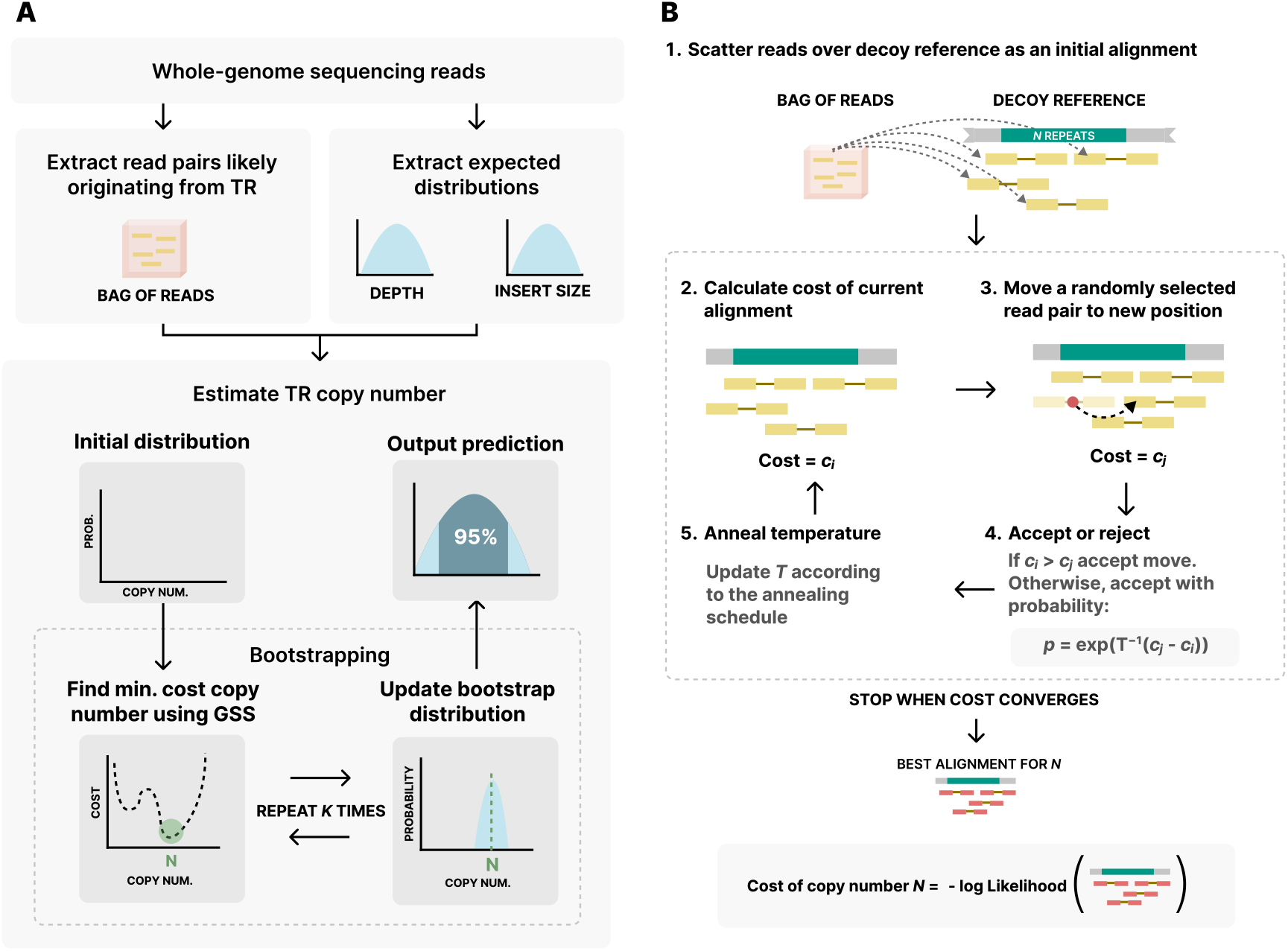
Overview of the ScatTR method. (A) Reads likely originating from the TR locus are collected from reference-aligned WGS data that includes both mapped and unmapped reads to form the “bag of reads”. Additionally, the expected read depth and insert size distributions are extracted. These distributions are used as parameters in the likelihood function. TR copy number estimation is done with bootstrapping. ScatTR iteratively finds the copy number with the minimum cost using golden section search (GSS) and uses the result to update the bootstrap distribution. Finally, the bootstrap distribution is used to output an estimate and a 95% confidence interval. (B) For a given copy number, ScatTR evaluates the cost by finding the best alignment to a decoy reference with the given number of repeat units. It starts with an initial alignment, then updates the alignment using Monte Carlo moves to reduce the cost and accept changes based on a probability function. This process is done via simulated annealing. It continues until convergence, yielding the best alignment for the given TR copy number. The best alignment is used to calculate the cost of the copy number, which is what GSS minimizes in (A).

While ScatTR is not designed to estimate the copy number of TRs shorter than the fragment length, we wanted to understand its accuracy relative to the state-of-the-art methods in such cases. Similar to our benchmarking for large TRs, we simulated heterozygous and homozygous expansions in 30 TR loci with copy numbers ranging from 5 to 45 for a total of 540 samples. We observe that ScatTR underperforms compared to other methods for TRs shorter than the fragment length, while ExpansionHunter consistently maintains the lowest RMSE across different copy numbers for these short TRs (Figure 2C).

**Figure 2:**
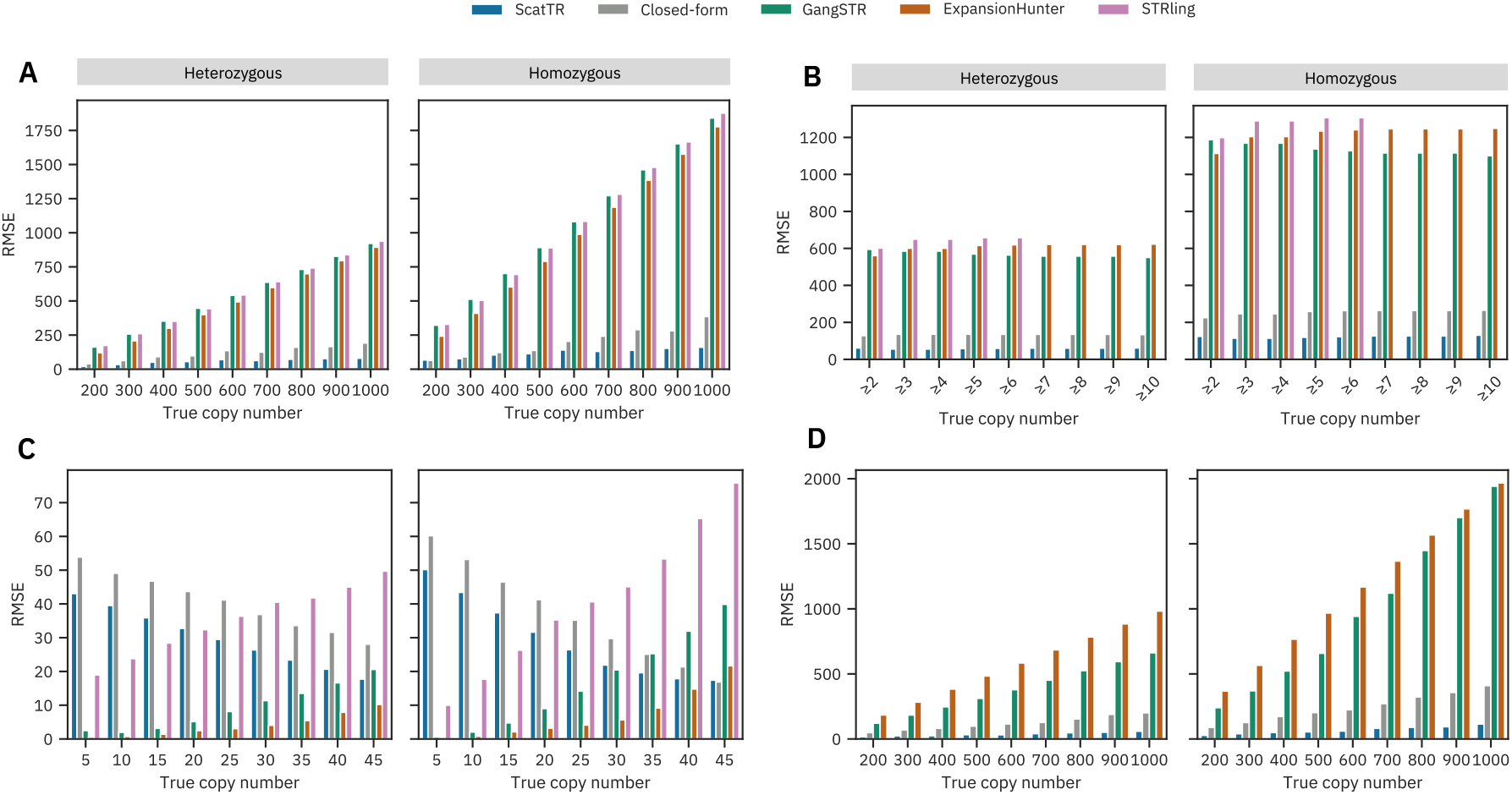
Benchmarking accuracy against the state-of-art with simulated data. We estimated the TR copy number using state-of-the-art methods (ExpansionHunter, GangSTR, and STRling), ScatTR, and a closed-form solution, and calculated the Root Mean Square Error (RMSE) between the predicted and the true copy number of TRs on simulated data. (A) RMSE across samples compared to true copy numbers for large TRs with motif lengths between 2-20 bp. Heterozygous expansions on the left, homozygous on the right. These estimates were conducted on 540 simulated short-read WGS samples representing 30 TR loci and a range of known copy numbers (200-1000). (B) RMSE as a function of TR motif length for (A). (C) RMSE across samples compared to true copy number of small TRs with motif lengths between 2-20 bp. Heterozygous expansions on the left, homozygous on the right. These estimates were conducted on 540 simulated short-read WGS samples representing 30 TR loci and a range of known copy numbers (5-45). (D) RMSE across samples compared to true copy numbers for large TRs with motif lengths between 21-50 bp. These estimates were conducted on 540 simulated WGS samples with TR expansions, representing 30 TR loci and a range of known copy numbers (200-1000).

Next, while ExpansionHunter and GangSTR were not specifically designed for large motifs, we wanted to test ScatTR’s performance on long motifs compared to these tools. STRling was excluded from this analysis since it does not allow users to input motifs larger than 6 bp. We sampled a subset of 30 TR loci with motif lengths 21-50 bp from TRs identified by TRF (Supplemental Table 2). We simulated homozygous and heterozygous expansions for the sampled TR loci with copy numbers ranging from 200 to 1000 for a total of 540 samples. We observe that ScatTR maintains the lowest RMSE across different TR copy numbers, with heterozygous TRs better predicted than homozygous TRs (Figure 2D).

### 2.3. ScatTR captures mismapped IRRs more effectively than existing methods

Accurately capturing and mapping IRRs is a key challenge in TR copy number estimation (Dolzhenko, Vugt, et al. 2017; Gymrek, Willems, et al. 2016; Dashnow, Pedersen, et al. 2022). IRRs originating from a TR of interest may be mismapped to other genomic regions or remain unmapped altogether. To assess ScatTR’s ability to recover these reads, we identified the ground truth IRRs in the 540 simulated samples described earlier in 2.2 and compared them to the IRRs extracted by ScatTR. Across all samples, ScatTR achieved a mean precision of 0.999 (SD = 0.003) and a mean recall of 0.954 (SD = 0.0635), indicating that it captures nearly all true IRRs, though a small fraction remains unretrieved or mismapped (Supplemental Figure 1).

Some existing tools, such as ExpansionHunter and GangSTR, attempt to improve IRR retrieval and mapping by incorporating user-defined off-target regions—genomic regions where IRRs are likely to mismap (Dolzhenko, Bennett, et al. 2020; Gymrek, Willems, et al. 2016). These methods scan predefined off-target regions to recover additional IRRs before estimating the TR copy number. In contrast, ScatTR scans the entire WGS alignment file (CRAM/BAM/SAM) without requiring predefined off-target regions, allowing it to adaptively identify relevant reads.

To evaluate how ScatTR performs relative to ExpansionHunter and GangSTR when off-target regions are used, we analyzed TR loci where each tools define such regions. Expansion-Hunter’s catalog includes *C9orf72* and *FMR1*, while GangSTR’s catalog contains 12 TR loci with off-target regions, including *SCA1–3, SCA6–8, SCA12, SCA17, HTT, DM1, FMR1*, and *C9orf72*. We simulated WGS samples with varying copy numbers for each locus and estimated TR copy numbers using ExpansionHunter and GangSTR both with and without off-target regions. As shown in Figure 3A, ScatTR’s RMSE was lower than 90.57 and 145.57 across all copy numbers for heterozygous and homozygous cases, respectively, while GangSTR’s RMSE was much higher with off-target regions seeming to slightly increase RMSE. Additionally, in Figure 3B, we see that off-target regions help decrease ExpansionHunter’s RMSE, but it is still much higher than ScatTR’s RMSE, which is less than 180.08 and 249.90 across all copy numbers for heterozygous and homozygous cases, respectively. ScatTR consistently outperforms ExpansionHunter and GangSTR, even when off-target regions were incorporated, demonstrating its superior ability to recover IRRs and accurately estimate TR copy numbers.

**Figure 3:**
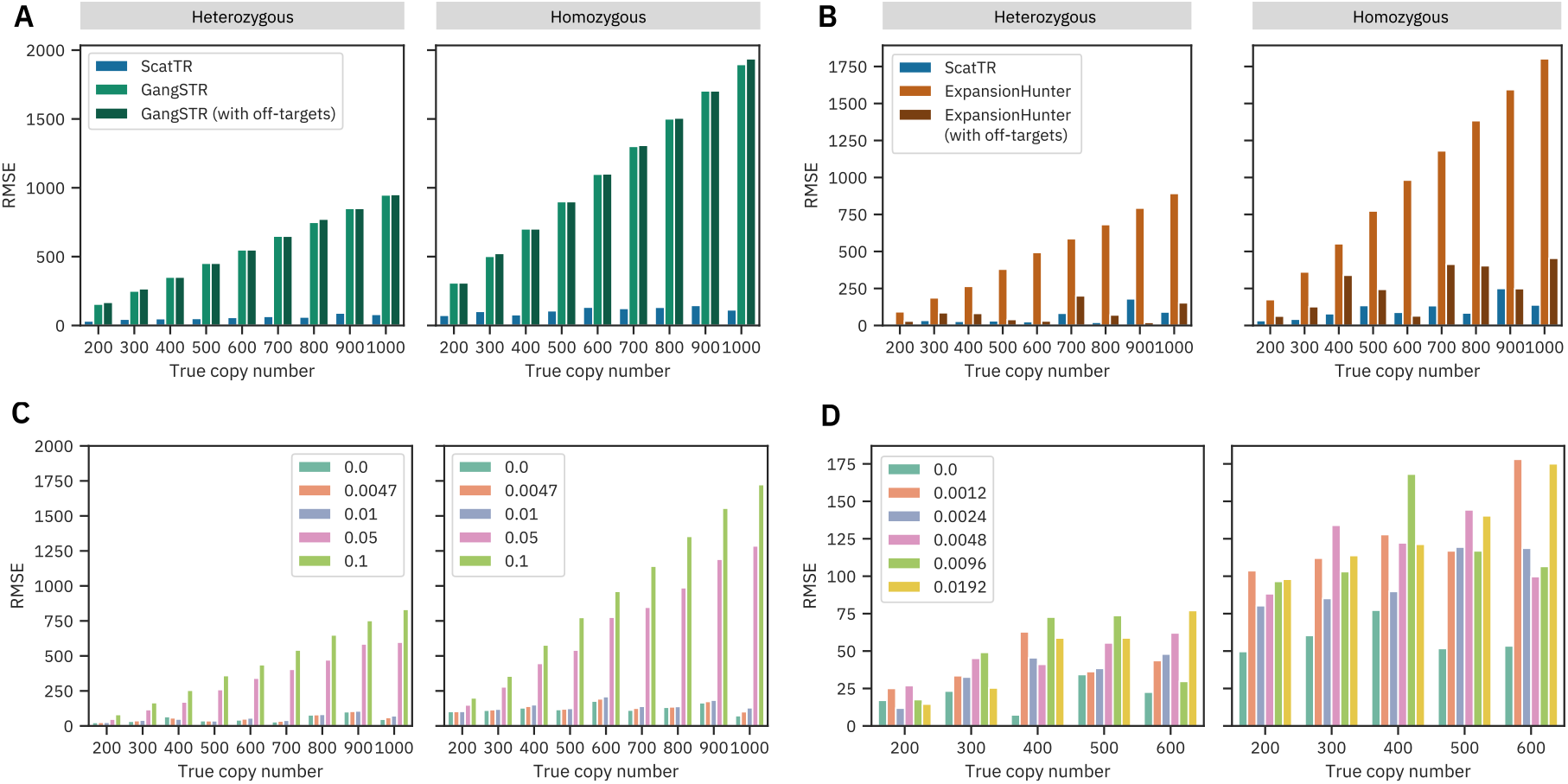
Impact of off-target regions, mutation rates, and sequencing errors on ScatTR accuracy. (A) RMSE of ScatTR compared to GangSTR, with and without off-target regions, in 108 simulated samples across different true copy numbers. Heterozygous cases are shown on the left, homozygous on the right. The simulated samples represent expansions in 12 TR loci for which GangSTR provided off-target regions, including *SCA1–3, SCA6–8, SCA12, SCA17, HTT, DM1, FMR1*, and *C9orf72*. (B) RMSE of ScatTR compared to Expansion-Hunter, with and without off-target regions, in 18 simulated samples across different true copy numbers. Heterozygous cases are shown on the left, homozygous on the right. The simulated samples represent expansions in 2 TR loci for which ExpansionHunter provided off-target regions, including *C9orf72* and *FMR1*. (C) RMSE of ScatTR across varying mutation rates in the repeat sequence. (D) RMSE of ScatTR across different sequencing error rates in whole-genome sequencing (WGS) samples.

### 2.4 ScatTR is robust against expected mutations and sequencing error

To evaluate ScatTR’s performance in the presence of mutations in the repeat sequence, we sampled 5 TR loci with motif lengths 2-20 bp (Supplemental Table 3). We simulated and aligned WGS samples with motif copy numbers ranging from 200 to 1000, both homozygous and heterozygous. For each locus, we systematically varied the proportion of mutated bases within the repeat sequence. As expected, the accuracy of the ScatTR improved as the mutation rate decreased (Figure 3C). We then calculated the expected mutation rate as 0.0047 at these loci using long-read sequencing data across five 1000 Genomes Project samples (Logsdon et al. 2024) (see 4.7). At the expected mutation rate, we found that the highest RMSE for ScatTR was 193.32 across all tested copy numbers, which is comparable to 177.59 when there are no mutations present. This indicates that our method effectively estimates TR copy numbers even in the presence of naturally occurring sequence variation.

To assess the effect of sequencing errors on ScatTR’s performance, we used the same five loci as above to simulate and align WGS samples with varying sequencing error rates. We generated reads using ART (Huang et al. 2012), a sequencing read simulator that models sequencing errors based on empirical error profiles from different sequencing platforms. It also allows for overriding the sequencing error rate with a user-specified value. The Illumina HiSeq X platform error profile, as provided by ART, assumes a sequencing error rate of approximately 0.0012. We simulated samples at error rates that were multiples of this baseline and measured ScatTR’s accuracy. As expected, ScatTR’s RMSE increased as the sequencing error rate increased (Figure 3D). However, at the baseline error rate of 0.0012, the highest RMSE was 62.82 and 178.11 across all tested copy numbers for heterozygous and homozygous cases, respectively, which is comparable to the RMSE of 34.33 and 77.28 when there are no sequencing errors.

### 2.5 Genome-wide profiling of large expanded TRs in the population

To demonstrate the utility of ScatTR, we used 2495 PCR-free high-coverage WGS data from the 1000 Genomes project (Byrska-Bishop et al. 2022). Since ScatTR and the other previous state-of-the-art methods require a catalog (except STRling), we first identified TR expansions larger than the read length with ExpansionHunter denovo (Dolzhenko, Bennett, et al. 2020), which detects novel repeats from WGS data and reports the number of fully IRR pairs. Since it only provides approximate coordinates, we overlapped the identified expanded TR regions with repeat regions identified by TRF to find their exact coordinates. We retained loci that had at least one fully in-repeat read pair in a minimum of 50 samples. ScatTR detected several large expansions in the 1000 Genomes population that go undetected by state-of-the-art methods (Figure 4A). Other methods consistently predict lengths shorter than or around the fragment length for the identified TR loci. In contrast, ScatTR estimates lengths larger than the fragment length (400 bp) for 6 out of 8 of the identified loci.

**Figure 4:**
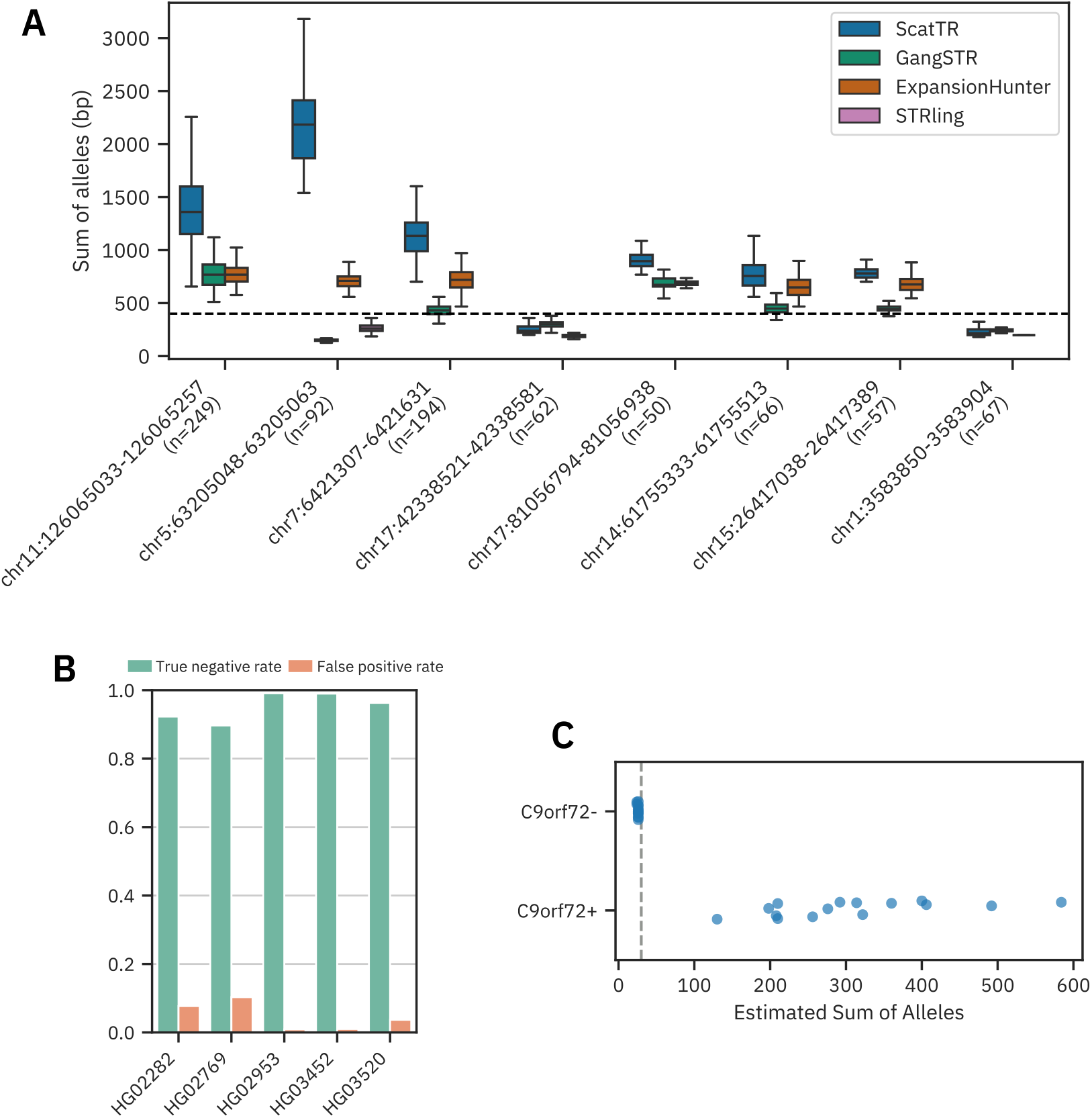
Evaluation of ScatTR on real WGS data. (A) Predicted sum of allele lengths (bp) of TR expansions by ScatTR and state-of-the-art methods. The labels on the x-axis indicate the sample size for each locus. All loci have motif lengths greater than 6 bp, except for the locus on chromosome 5. The mean fragment length of 400 bp is shown as a horizontal line. (B) True negative rate and false positive rate of TR expansion calls for five samples with long-read ground truth across 171146 TR loci. Expansions are defined as exceeding the fragment length threshold of 400 bp. (C) Predicted sum of allele lengths (bp) for 30 ALS patients based on short-read WGS data. The dataset includes 15 samples with known C9orf72 expansions (C9orf72+) and 15 without (C9orf72-). The vertical line indicates the pathogenic cutoff of 30 repeat units. All samples were correctly classified.

However, because of the concern of fully IRR pairs originating from other TR loci contaminating the bag of reads, we wanted to assess the false positive rate of expansions larger than the fragment length called by ScatTR. We used five 1000 Genomes Project samples (Logsdon et al. 2024) that have both long-read and short-read WGS available. We first applied TRGT (Dolzhenko, English, et al. 2023) to the long-read data to calculate the length of 171,146 TR loci provided by TRGT catalogue (see 4.7). We then used ScatTR on short-read data to estimate the length of the same TR loci. When the long-read data is used as gold standard, we found that, as shown in Figure 4B, the mean false positive rate for ScatTR is extremely low, 0.047.

### 2.6 ScatTR correctly identifies samples with C9orf72 pathogenic repeat copy numbers

To further validate the utility of ScatTR, we assessed its ability to estimate the size of a well-characterized pathogenic repeat expansion in *C9orf72*, a biomarker for familial ALS and frontotemporal dementia (FTD). We analyzed short-read WGS data from 30 ALS patients (Kenna et al. 2016), including 15 individuals with a known expansion (*C9orf72* +) and 15 without (C9orf72-). As shown in Figure 4C, ScatTR effectively distinguished *C9orf72* + patients from *C9orf72* - patients with 100% accuracy. This demonstrates that ScatTR can reliably identify large pathogenic repeat expansions from short-read data, a key challenge in the study of repeat expansion disorders.

### 2.7 Algorithm resource requirements

To evaluate ScatTR’s runtime performance, we benchmarked its execution time against state-of-the-art methods on five whole-genome sequencing (WGS) samples from the 1000 Genomes Project (Byrska-Bishop et al. 2022): HG03520, HG02282, HG02953, HG02769, and HG03452. Each tool was tested on randomly subsampled TR catalogs of varying sizes (1, 10, 100, 1000, and 10,000 loci) from the TRF catalog. All benchmarks were conducted on a Dell PowerEdge R6625 with 2 x AMD EPYC 9254 24-Core Processors, operating in single-threaded mode. Memory usage was negligible across all methods.

As shown in Figure 5A, ScatTR required an average of 27.66 minutes per sample to estimate the copy number for one locus. Alternative tools such as GangSTR and ExpansionHunter completed the task significantly faster, while STRling performed more efficiently than ScatTR, requiring about half the execution time. However, the majority of ScatTR’s runtime—an average of 26.94 minutes—was spent on read extraction.

**Figure 5:**
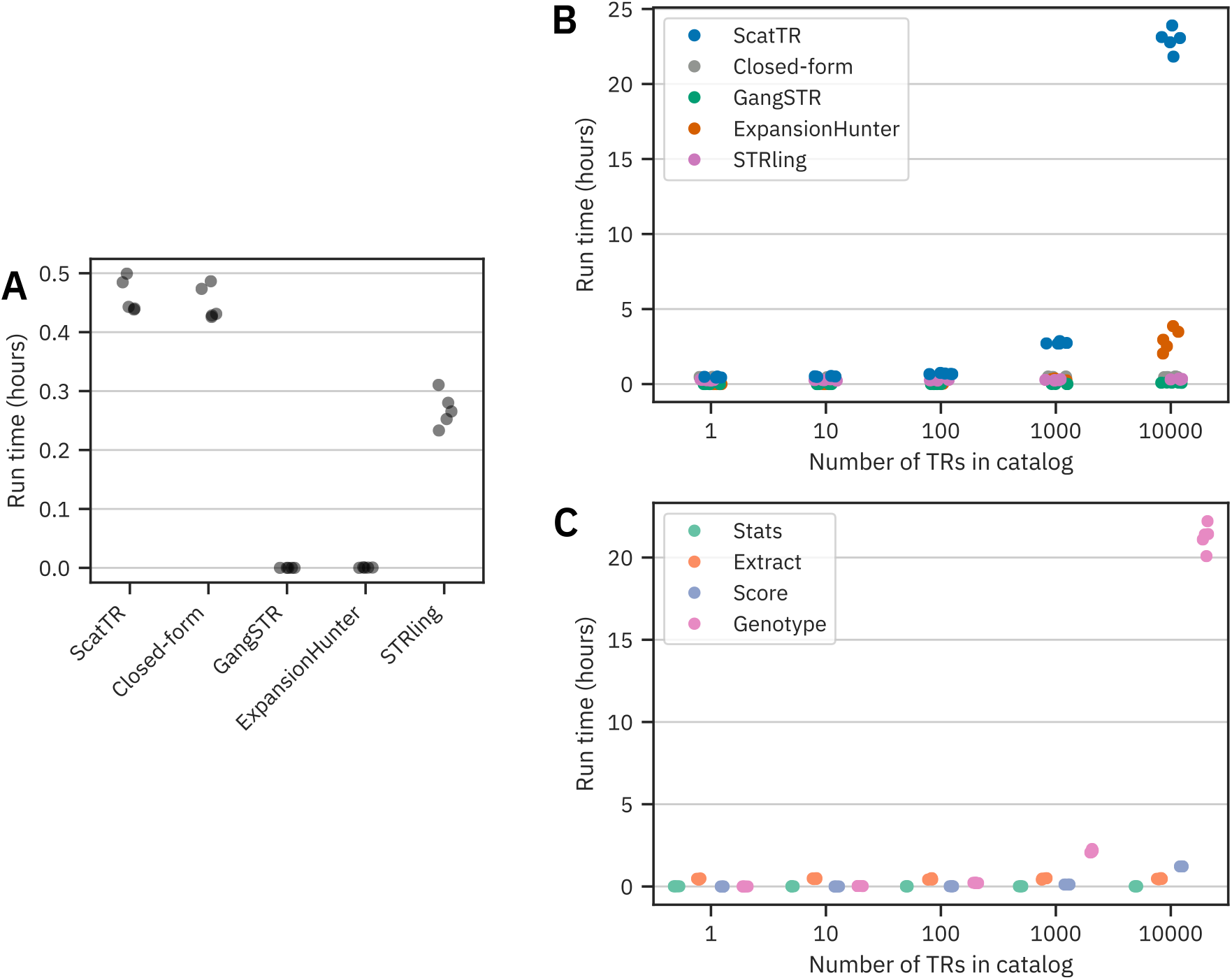
Runtime evaluation of ScatTR in simulated and real WGS samples. (A) Measured runtime (hours) for estimating TR copy number at a single locus across 5 real WGS samples. Reported times are the mean of five runs. (B) Measured runtime (hours) for estimating TR copy numbers across varying catalog sizes in 5 WGS samples, comparing ScatTR, state-of-the-art methods, and a closed-form solution. (C) Breakdown of ScatTR runtime by processing step for (B).

As TR catalog size increased, so did the runtime for all methods. For ScatTR, the read extraction and expected distribution extraction steps remained constant, averaging 25.20 minutes and 0.57 minutes per sample, respectively. However, the read scoring step (which calculates alignment scores for reads in the bag against decoy references, see 4.1.9) and the genotyping step (which involves multiple bootstrap runs of GSS and Monte Carlo simulations, see 4.1.7) varied with catalog size. At the largest catalog size (10,000 loci), ScatTR required an average of 22.94 hours per sample, while the next slowest tool, ExpansionHunter, averaged 2.97 hours (Figure 5B). A breakdown of ScatTR’s runtime by processing step (Figure 5C) shows that most of the time was spent on scoring and genotyping. On average, ScatTR processed one locus in 8.08 seconds, compared to ExpansionHunter’s 1.06 seconds per locus. This trade-off reflects ScatTR’s focus on accuracy for large TRs.

## 3. Discussion

We present ScatTR, a novel method for estimating the copy number of TRs that exceed the fragment length in paired-end short-read WGS data. ScatTR introduces an innovative approach for estimating TR copy numbers. Unlike prior methods that classify reads into distinct categories such as flanking, enclosing, spanning, or anchored IRR pairs (Gymrek, Golan, et al. 2012; Willems et al. 2017; Dashnow, Lek, et al. 2018; Tang et al. 2017; Tankard et al. 2018; Dolzhenko, Vugt, et al. 2017; Mousavi et al. 2019), ScatTR uses a more flexible and comprehensive approach, overcoming key limitations of these classification-based strategies.

Existing methods typically exclude fully IRR pairs from copy number estimation unless the user specifies off-target regions where these pairs may have been incorrectly mapped. Our approach integrates fully IRR pairs directly into the estimation process. The closed-form solution, which incorporates fully IRR pairs, demonstrates significant improvements in accuracy for large TRs. However, incorporating these reads alone is insufficient for achieving high accuracy, necessitating the development of a more flexible likelihood function to further refine copy number estimates.

Unlike prior approaches where the likelihood is determined by the number of read pairs in each category and a category-specific characteristic, ScatTR adopts a likelihood model inspired by genome assembly methods. In genome assembly, the likelihood of an assembly is based on all read alignment combinations to the assumed true genome assembly (or reference). Recent methods approximate this likelihood by only using a few best read alignment combinations acquired using traditional read alignment methods such as BWA (Boža et al. 2015). However, ScatTR approximates the likelihood for a copy number using only the best alignment. Finding the best alignment to a decoy reference (containing a large repetitive region) is an ambiguous task since reads from within the repeat can perfectly align to multiple locations along the decoy reference. Thus, traditional read alignment methods are not helpful for finding the best alignment in TR copy number determination. This is also why it is difficult to assemble repetitive parts of genomes using short reads (Treangen and Salzberg 2012). ScatTR overcomes this challenge by using Monte Carlo sampling to efficiently find the best alignment to decoy references. It incorporates information from both insert size and depth distribution, the latter of which is not used in genome assembly likelihood models, since that information is not known for that task.

Our results show that this novel approach enables accurate estimation of TR lengths that exceed typical fragment lengths. ScatTR outperforms state-of-the-art methods, particularly for larger TRs and longer motifs up to 50 bp. Unlike classification-based approaches, ScatTR’s model does not rely on directly counting repeat units within read sequences, allowing it to accommodate longer motifs with greater flexibility. Additionally, ScatTR offers the added advantage of generating a plausible alignment of reads to the predicted TR copy number. This alignment capability could facilitate deeper analysis of repeat mosaicism and interruptions. However, further testing is required to fully assess the utility of these alignment solutions.

Although ScatTR is computationally more demanding than existing approaches, this tradeoff is a direct consequence of its design to maximize accuracy for large TRs. Unlike methods that optimize efficiency by analyzing only a subset of reads associated with a TR, ScatTR processes all available reads spanning the repeat region, including in-repeat reads that may be unmapped or mismapped elsewhere in the genome. This requires scanning the entire referenced-aligned NGS data including both mapped and unmapped reads, which increases computational load. Additionally, ScatTR’s Monte Carlo approach improves accuracy beyond that of closed-form solutions based solely on IRR read counts. This exhaustive strategy is particularly valuable for large pathogenic TR expansions, where precise repeat length determination might be critical for clinical interpretations. Given that no current method accurately genotypes very large TR expansions using short-read sequencing, we believe the trade-off between computational efficiency and accuracy is well justified, particularly for targeted analyses of TR loci.

While ScatTR excels in estimating simple TRs, it does not currently handle complex repeat structures that consist of multiple motifs or known interruptions. This limitation arises from the efficiency optimizations in ScatTR that assume a simple repeat structure. Notably, tools like STRling and GangSTR also do not support complex loci, whereas ExpansionHunter and Dante can handle such loci for short TRs, provided that the repeat structure is accurately defined. Future research could focus on algorithmic enhancements to extend ScatTR’s applicability to complex loci, an important category of TR variation with significant implications for human genetic studies and disease research.

## 4. Methods

### 4.1. The ScatTR algorithm

In this section, we detail the steps of the ScatTR algorithm as shown in Figure 1. In 4.1.1, we describe how the expected distributions are extracted, and in 4.1.2 how the bag of reads is extracted—both of which are used as inputs to the copy number estimation process.

The estimation process relies on optimizing the likelihood function which we derive in 4.1.3. To compute the likelihood of a copy number, the best alignment of the bag of reads needs to be found. We describe a Monte Carlo process in 4.1.7 to find the best alignment. Then, to efficiently search the space of all copy numbers to find the one with the highest likelihood value, we describe an optimization process in Section 4.1.8. Finally, we explain how we simulated WGS data to test the performance of our method and how we compared it to various methods developed previously.

#### 4.1.1. Expected depth and insert size distributions

To extract the expected read depth distribution from WGS short-read data, we sample aligned reads from 100 random regions, excluding those with average mapping quality *≤* 60. Each position’s depth within the sampled regions contributes to the distribution. Then, the distribution is trimmed to the 99th percentile to remove outlier depth values.

To extract the expected insert size distribution from WGS short-read data, we sample aligned reads from 100 random regions. A read’s observed insert size, as determined by the aligner, is included in the distribution if the read meets specific criteria: it must be a primary read, part of a proper pair, unclipped, and have a mapping quality of at least 60.

#### 4.1.2 Extracting the bag of reads

We extract the paired-end reads that are likely to originate from the TR locus and refer to them as the “bag of reads”. These reads fall into three categories: anchored IRR pairs, fully IRR read pairs, and spanning pairs.

Anchored IRR pairs have one mate that lies within the repeat region and the other lies on either the left or right flanking region of the repeat (thus ‘anchoring’ the pair). The flanking regions are the regions outside the repeat. Anchored IRR pairs only occur if the repeat is larger than the read length. Otherwise, it would be impossible to observe the mate that lies within the repeat region.

Fully IRR pairs occur only when the repeat is larger than the fragment length. Otherwise, we would never observe such pair. Previous methods do not incorporate information from fully IRR pairs since they are not ‘anchored’ to the flanking region of the repeat and therefore cannot be unambiguously assigned to the locus. An advantage of our algorithm is that it relies on an optimization criterion that decides whether a fully IRR pair belongs to (*i*.*e*., can be aligned to) a specific locus (see Section 4.1.7).

Spanning pairs are those in which both mates map entirely within the flanking regions, surrounding but not overlapping the repeat itself. These pairs provide additional information about the repeat’s boundaries and are included in our analysis.

To extract these read pairs, we scan the alignment file (BAM/CRAM/SAM) for reads with at least one mate mapped to the flanking regions of the repeat (a window of read-length size around the reference repeat region). This process captures both anchored IRR pairs and spanning pairs. Fully IRR pairs are identified separately by testing all read pairs with mapping quality *≤*40. If both mates in a pair are classified as IRRs, the pair is also added to the bag of reads.

A read is classified as an IRR if its sequence matches with the expected repeat pattern closely, even after being rotated or reverse complemented. Dolzhenko et al. (Dolzhenko, Vugt, et al. 2017) describe a weighted purity (WP) score to quantify how closely a read matches the expected repeat pattern. To compute the WP score, we compare the read to the nearest perfect repeat sequence across these transformations (*e*.*g*., a CAG repeat can manifest as CAG, AGC, GCA in the forward direction or as CTG, TGC, GCT in the reverse direction). The WP score accounts for both matching bases and mismatches, with scores of 1 for matches, 0.5 for low-quality mismatches, and −1 for high-quality mismatches. The resulting score is normalized by the read length to produce a WP value between −1 and 1. Reads with a WP score above the threshold are deemed likely to originate entirely from the repeat region. Here we used a threshold of 0.9 (default in our tool).

#### 4.1.3 Likelihood of a copy number

To determine the most probable TR copy number from a given bag of reads (i.e., set of sequencing reads), we derive the likelihood function that quantifies how well different candidate copy numbers explain the observed data. This process involves defining the probability of observing a set of reads given a proposed copy number and an alignment of those reads to a reference sequence (which we call a decoy reference). In this section, we introduce a general probability model that accounts for all possible alignments of the reads. Next, we refine this model by identifying the best alignment, which allows us to approximate the likelihood function more efficiently. By breaking down the likelihood into its component probabilities—in Section 4.1.4 and Section 4.1.5–we establish a structured framework for computing the likelihood in a way that is both theoretically sound and computationally feasible. Finally, we put together these building blocks in Section 4.1.6 to define the full form of the likelihood that ScatTR uses and adapt it to account for diploid genomes, ensuring our model is applicable to real biological data.

Here, we will derive Pr(*R* | *C*), the probability (or likelihood) that a bag of reads *R* is observed assuming that *C* is the true TR copy number from which *R* was sequenced. In the ScatTR algorithm, we use this likelihood to compare proposed copy numbers for a given bag of reads with the goal of finding the copy number that maximizes the likelihood. The TR copy number that maximizes the likelihood is the most probable TR copy number given the bag of reads.

To show that maximizing the likelihood, Pr(*R* | *C*), is equivalent to maximizing the probability of the TR copy number given the bag of reads, Pr(*C* | *R*), we apply Bayes’ rule:

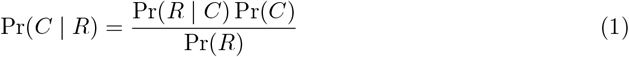

In the equation, Pr(*C*) is the prior probability of the TR copy number. We assume that this prior probability is constant across the set of reasonable copy numbers for a given *R* (see 4.1.8 for how a reasonable set is calculated). Pr(*R*) is the prior probability of observing the bag of reads. Since our primary goal is to compare the likelihood of various copy numbers for the same bag of reads, we can assume Pr(*R*) is a constant. Therefore, for the purpose of comparing TR copy numbers, maximizing the values Pr(*C* | *R*) and Pr(*R* | *C*) over *C* is equivalent.

Previous work has typically modeled the likelihood Pr(*R* | *C*) by categorizing reads into distinct classes like flanking, enclosing, spanning, or anchored in-repeat read pairs (Gymrek, Golan, et al. 2012; Willems et al. 2017; Dashnow, Lek, et al. 2018; Tang et al. 2017; Tankard et al. 2018; Dolzhenko, Vugt, et al. 2017; Mousavi et al. 2019). In these studies, the likelihood is determined by both the count of read pairs in each class and a characteristic specific to that class. In contrast, we do not define the likelihood by classifying reads into classes and modeling their characteristics. Instead, we use an approach similar to the one used in genome assembly problems, where the likelihood is based on the alignments of the observed reads to the given genome assembly (i.e., TR copy number in our case).

The problem of finding the TR copy number that maximizes the likelihood of observing a set of reads can be reduced to finding the genome assembly that maximizes this likelihood. Because in genome assembly, the TR copy number is determined as part of constructing the most likely genome sequence given the reads. When we search for the genome assembly that maximizes the likelihood of observing the given set of reads, we consider all possible copy numbers. Therefore, finding the assembly that best explains the reads directly provides the copy numbers that maximize the likelihood. Thus, the problem of finding the correct TR copy number reduces to finding the correct genome assembly.

There exist several definitions of the likelihood of reads given a genome assembly (Rahman and Pachter 2013; Clark et al. 2013; Ghodsi et al. 2013). Previous studies have shown that the likelihood is typically maximized for correct genome assemblies. Therefore, we can adapt these likelihood functions with the expectation that finding their maximum will result in finding the correct TR copy numbers.

However, several adaptations need to be made. Based on Ghodsi et al. (Ghodsi et al. 2013), we can define the likelihood Pr(*R* | *C*) by considering all possible alignment positions of the reads in *R* to the decoy reference *D*_*C*_. The decoy reference *D*_*C*_ is constructed by concatenating the left flanking region of the repeat, *C* repeat units, and the right flanking region of the motif. Note that in the context of finding the TR copy number that maximizes the likelihood of the reads, we are constructing multiple decoy references with varying values of *C*.

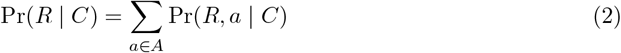

where *A* is the set of all alignments of the reads. A single alignment *a* is a mapping of each read to its positions and orientation (forward or reverse) along the decoy reference *D*_*C*_. The alignment refers to the positions only and does not include pairwise alignment information. To compute the exact value of Pr(*R* | *C*), Ghodsi et al. developed a dynamic programming algorithm that marginalizes over all possible alignments of the reads in *R* to the reference. However, this algorithm is resource-intensive, and is mainly used to compute the quality of assembled genomes, and not to find a high likelihood assembly. In practice, only several best alignments contribute significantly to the overall probability (Boža et al. 2015). We approximate the likelihood with the best alignment 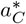of the reads to decoy reference 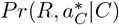is used to approximate the overall likelihood *Pr*(*R*|*C*) because it contributes the largest value. In 4.1.7, we describe how we find this best alignment. This likelihood is defined as:

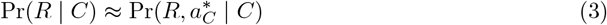

Now, we define *P* (*R, a* | *C*), the probability of a fixed alignment *a* of *R* given the copy number *C*. Using the chain rule, the joint probability *P* (*R, a* | *G*) can be expressed as the product of a conditional probability and a marginal probability:

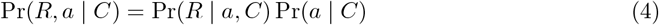

where Pr(*R* | *a, C*) is the probability of the reads given that the true alignment is *a* and the TR copy number is *C*, and 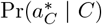is the probability of the alignment positions given that the true copy number is *C*. In sections 4.1.4 and 4.1.5, we define the two probability terms that make up Pr(*R, a* | *C*) in general form, independent of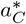. This allows us to model alignment probability in a general sense before incorporating the best alignment in 4.1.6.

#### 4.1.4. Probability of observing a set of reads given their true alignment and TR copy number

The model assumes that individual reads are independently sampled, thus the overall probability Pr(*R* | *a, C*) is the product of the individual read mapping probabilities. To model the read mapping probability, we incorporate a simple sequencing model that accounts for substitution errors (Boža et al. 2015). Then, the probability for a single read *r* aligned to position *j* is:

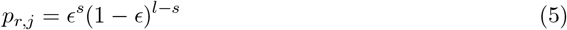

where *∈*is the sequencing error rate, *s* is the number of substitutions relative to *D*_*C*_ at position *j* and *l* is the length of the read *r*

Then, the overall probability becomes:

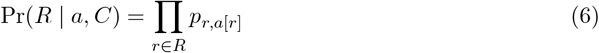

where *a*[*r*] is the position of *r* in the alignment *a*.

#### 4.1.5. Probability of observing an alignment given the true TR copy number

The alignment *a*, regardless of the exact sequences of the reads, correspond to an observed insert size and read depth distribution. The read depth is the number of reads that overlap a position along the decoy reference *D*_*C*_. We expect both these observed distributions to be close to the expected distributions of the sample (see 4.1.1).

We model the probability of the alignment *a* given the true TR copy number, Pr(*a* | *C*), as the product of the observed insert size and depth distribution probabilities.

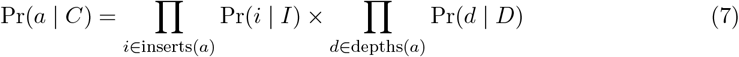

where *I* and *D* are the expected insert size and read depth distributions of the sample, respectively. inserts(*a*) and depths(*a*) are the sets of insert sizes and read depths implied by the alignment, respectively.

#### 4.1.6 Overall likelihood of the bag of reads given a true TR copy number for a diploid sample

We combine the equations 3, 6 and 7 into the the overall approximate definition of the likelihood:

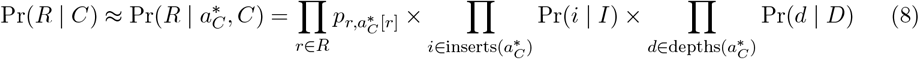

where 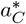is the best alignment of *R* to the decoy reference *D*_*C*_.

This derivation assumes a single haplotype. In practice, human genomes are diploid and may have different copy numbers (also known as alleles) on each haplotype. To overcome this, ScatTR aligns reads to two decoy haplotypes (maternal and paternal) and assumes that read depth is evenly distributed between them. To adapt our derivation, let *C*^*′*^= *⟨C*_1_, *C*_2_*⟩*, where *C*_1_ and *C*_2_ are the corresponding copy numbers on each haplotype. We redefine the likelihood as such:

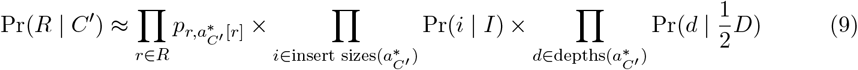

The alignment 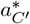is now a mapping of each read to its position, orientation, and additionally the haplotype. A read can only be aligned to one haplotype. An alignment is only valid if all pairs are aligned to the same haplotype and in opposing orientations. The likelihood is not defined for invalid alignments where pairs are aligned to different haplotypes or have the same orientation. 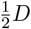 represents the halved depth distribution, assuming that reads are equally likely to originate from either haplotype.

#### 4.1.7 Finding the best alignment for a copy number

Here we describe a Monte Carlo optimization routine that finds the best alignment 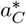of reads *R* to a decoy reference *D*_*C*_ among all possible alignments *A*_*C*_. The alignment is a mapping of each read to its position and orientation on the decoy reference. In 4.1.3, we describe how this best alignment is used to approximate the likelihood of *R* given the true copy number. The likelihood of the alignment Pr(*R, a* | *C*) is used to approximate the overall likelihood Pr(*R* | *C*) because it contributes the largest value to the overall likelihood. So, we seek to find:

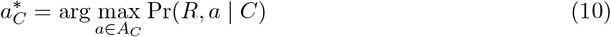

Finding 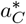is difficult because the search space, *A*_*C*_, is combinatorial in size with respect to *C* and |*R*|. Therefore, we use simulated annealing (SA) which is an iterative probabilistic technique for approximating the global minimum of a cost function (Kirkpatrick et al. 1983). It is especially useful when the search space is large, and often used when the search space is discrete.

In our implementation, we seek to minimize the following cost function, which is the negative log likelihood of observing the reads *R* given the true alignment *a* and copy number *C*. We derived the likelihood in 4.1.3.

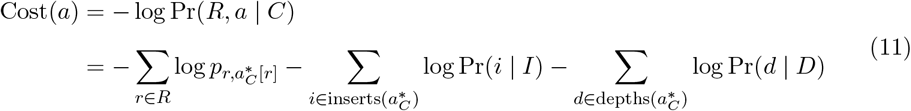

We start from a randomly initialized alignment *a* and an initial temperature *t*_0_ (a parameter of the SA algorithm). In each iteration *i*, we choose a number of read pairs to move, and randomly sample new positions on the decoy reference to align them to. This forms a new alignment *a*^*′*^. The number of read pairs that are moved is proportional to the temperature *t*_*i*_. The next step depends on the costs of the alignments *a* and *a*^*′*^:

1. If Cost(*a*^*′*^) *≤* Cost(*a*), then the new alignment *a*^*′*^ is accepted and the algorithm continues with the new alignment
2. If Cost(*a*^*′*^) *>* Cost(*a*), then the new alignment *a*^*′*^ is accepted with probability proportional to the difference in cost. If *a*^*′*^ is rejected, the algorithm keeps the old alignment *a* for the next step

The acceptance probability is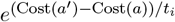. When the temperature *t*_*i*_ is high, changes to the alignment that increase the cost substantially (decrease the likelihood) are more likely to be accepted. We use a standard cooling schedule where 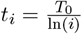at iteration *i* of the algorithm. The algorithm stops when no new best solutions are found for 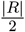 ln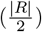iterations where is the number of read pairs. If no new best solutions are found for 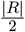iterations, the algorithm resets the temperature to the initial temperature *t*_0_. When the algorithm terminates, the alignment with the lowest observed cost is considered the optimal alignment, denoted as 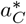.

#### 4.1.8 Finding the copy number that maximizes the likelihood of the bag of reads

To find the most likely copy number *C*^*^, we find the copy number that maximizes the likelihood of the observed bag of reads Pr(*R* | *C*). We explain in 4.1.3 how maximizing the former is equivalent to the later, and show how its value can be approximated using the best alignment for the given *C*. Thus, our goal is to find *C*^*^

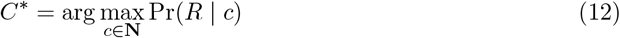

Every evaluation of Pr(*R* | *c*) requires finding the corresponding best alignment 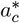,which is used to calculate the value of Pr(*R* | *c*). Since the search space is the set of natural numbers, it is inefficient to compute the likelihood for every copy number to find the maximum likelihood. Optimization techniques such as gradient descent are not appropriate since the likelihood function is non-differentiable.

An applicable technique is golden section search (GSS) (Kiefer 1953). It is an iterative optimization technique used to find a local extremum of a unimodal function within a specified interval. It works by successively narrowing the interval where the extremum lies, reducing the search space in each iteration. It significantly reduces the number of evaluations that are needed to find the copy number that maximizes the likelihood. Since it requires defining an initial interval, we calculate a reasonable range of copy numbers using a closed-form solution that relies on the mean depth of the sample and the number of in-repeat reads present in the bag. For the minimum value of the range we compute the closed-form estimate using a depth value of *µ*_depth_ + 3 _depth_, and the maximum using *µ*_depth_ *-* 3 _depth_. The closed-form estimate takes as input the number of IRRs observed and the expected mean depth (see 4.3).

In practice, to maintain numerical stability, we use the negative log likelihood as the minimization objective of GSS. Additionally, because GSS is designed to converge to any local maximum within an interval, it can sometimes converge to saddle points in our likelihood function. To improve our estimate of *C*^*^, we use bootstrapping to perform GSS multiple times. We start with an initial bootstrap distribution with the probability set to 0 for all copy numbers. For every bootstrap iteration, the copy number returned by GSS gets its probability incremented by the inverse of its the negative log likelihood. After bootstrapping is finished, the bootstrap distribution is normalized to sum to one and the median is reported as the estimate. This allows us to report a 95% confidence interval.

#### 4.1.9 Efficiently scoring reads against repeat decoy references

When trying to find the best alignment for a set of reads to a decoy reference genome, we need to efficiently compute a set of permitted alignment positions for each read and the associated alignment error. By default, we compute the alignment error as the Hamming distance between the read sequence and the corresponding reference sequence, separately for both the forward and reverse directions. Additionally, we provide an option for users to compute the alignment error using the Levenshtein distance, which accounts for insertions and deletions. This flexibility allows users to choose the appropriate metric based on the characteristics of the locus being analyzed, particularly for loci where indels are expected to play a significant role.

For a given read *r*, we first define “left flank” as the positions relative to the TR start position, “right flank” as the positions relative to the TR end position, and “within the TR” corresponds to the positions along the repeat motif. We only permit alignment positions with the smallest error, which could be more than one position. We store these positions in relative coordinates (relative to start or end of the TR) to help us generate positions in absolute coordinates for arbitrary length reference genomes. This means that we only need to compute the error at *F*_*L*_ + |*r*| *-* 1 positions for the left flank, *F*_*R*_ + |*r*| *-* 1 for the right flank, and *M* positions within the motif where *F*_*L*_ is the left flank length, *F*_*R*_ is the right flank length, and *M* is the repeat motif length. To generate the set of permitted alignment positions for an arbitrary reference (i.e., arbitrary TR copy number), we convert the precomputed relative positions to absolute positions on the reference. For relative positions on the left flank, we simply add *F*_*L*_. For relative positions on the right flank, we add *F*_*L*_ + *R*, where *R* is the length of the repeat region. For relative position *i* along the repeat motif, we generate all absolute positions *j* such that *j* = *F*_*L*_ + *n * i* for *n 2* **N** and *j* + |*r*| *≤ F*_*L*_ + *R*. This follows from the periodic nature of the TR sequence.

### 4.2. Simulating samples with tandem repeat expansions

We obtained a set of tandem repeat loci from the Simple Repeats catalog (hg38 coordinates) published on the UCSC genome browser (Kent et al. 2002). The catalog was generated by Tandem Repeats Finder (Benson 1999). First, we removed loci with a motif length shorter than 2bp, longer than 20bp. Additionally, we selected for loci that perfectly lift over to the T2T-v2 assembly (Nurk et al. 2022; Kent et al. 2002). A locus passes the filter only if the region *±*550 bp around the repeat locus is identical in both HG38 and the T2T-v2 assembly. We then randomly selected 30 loci for evaluating the performance of our method (Supplemental Table 1).

We used the T2T-v2 assembly (Nurk et al. 2022) as the reference genome from which WGS with paired short-read sequencing were simulated. We used ART (Huang et al. 2012) which simulates reads with customized error profiles. We generated our samples with the HiSeqX PCR-free profile, with 30x coverage, 150 bp read length, 450 bp mean fragment size, and 50 bp fragment size standard deviation. All other parameters were kept to the default. Specifically, the substitution error rate is approximately 0.0012. The insertion error rates are 0.00009 and 0.00015 for the first and second reads, respectively. The deletion error rates are 0.00011 and 0.00023, respectively. For each locus, we simulated a set of samples corresponding to copy numbers of 200 to 1000, with a step size of 100, as well as heterozygous and homozygous status. So, for each locus, we simulated 18 samples. In total we simulated 540 samples.

### 4.3. Baseline closed-form solution

As a baseline, we derived a closed-form solution to predict the copy number. The solution is equivalent to the copy number that maximizes the likelihood of the number of in-repeat reads (IRRs) observed. We counted a read as an IRR if it has a weighted-purity score *2* 0.9 (see 4.1.2).

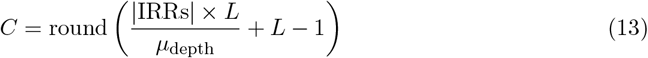

where *L* is the read length and *µ*_depth_ is the mean read depth of the sample. In the diploid case, we multiply the above value by 2 to get the sum of the copy numbers on both haplotypes.

### 4.4. Comparison with GangSTR

We obtained the source code for GangSTR version 2.5.0 from its Github repository (https://github.com/gymreklab/GangSTR). The binary was built using `htslib-1.11` bindings. We ran the simulated samples with the `--targeted` option. Since we are only interested in the genotype and estimated length, we used `--skip-qscore`. The `--readlength` option was set to `150`, `--coverage` to `30`, `--insertmean` to `300`, `--insertsdev` to `50`. We also set `--bam-samps` to the name of the sample and `--samp-sex` to `M` since chromosome Y was included in the reference from which reads were simulated from. The predicted repeat copy number is extracted from the output JSON file selected as the allele with the least absolute error to the true copy number.

### 4.5 Comparison with ExpansionHunter

We obtained the source code for ExpansionHunter version 5.0.0 (https://github.com/Illumina/ExpansionHunter). We ran the simulated samples using default parameters, and set the `--sex` option to `male`. The predicted repeat copy number is extracted from the output JSON file selected as the allele with the least absolute error to the true copy number.

### 4.6. Comparison with STRling

We obtained the source code for STRling version 0.5.2 (https://github.com/quinlan-lab/STRling). We ran the simulated samples using default parameters. Since STRling is not capable of analyzing TRs with motif lengths larger than 6, samples with such motifs were excluded from STRling predictions.

### 4.7. Calculating the expected mutation rate of TRs

We downloaded long-read sequencing data for HG02282, HG02769, HG02953, HG03452, and HG03520 from the 1000 Genomes Project portal Logsdon et al. 2024. Reads were aligned to the GRCh38 reference genome using `pbmm2 align --sort -j 48 <reference> <reads> <output>` using version 1.13.1. TR loci were then genotyped with TRGT (version 1.5.0) Dolzhenko, En-2023glish, et al. using `trgt genotype -t 6 --genome <reference> --repeats <repeats>--reads <reads> --output-prefix <output>`. From the output VCF file, we extracted the `AP` field from the `FORMAT` column for each locus, which represents the allele purity score. We computed the average purity across all samples and subtracted it from one to estimate the mean TR mutation rate.

## 5. Code availability

ScatTR is an open-source software package available for download at GitHub (https://github.com/g2lab/scattr). The source code, installation instructions, and documentation are provided in the repository. The software is implemented in Rust and supports both Linux and Mac operating systems.

## 6. Data access

PCR-free WGS data from participants with CANVAS are available from the Sequence Read Archive (SRA) under project PRJNA885420, with individual SRA accessions SRR21753324-SRR21753328. The dataset can be accessed at https://www.ncbi.nlm.nih.gov/sra.

PacBio HiFi long-read sequencing data were obtained from the International Genome Sample Resource (IGSR) data portal for the following samples: HG03520, HG02282, HG02953, HG02769, and HG03452.

The individual-level data from the NYGC ALS Consortium are available to authorized investigators through the dbGaP via accession code phs003067.

Additionally, PCR-free short-read WGS data from the 1000 Genomes Project are publicly available from the NIH 1000 Genomes mirror at https://ftp-trace.ncbi.nih.gov/1000genomes/ftp/1000G_2504_high_coverage/.

## Acknowledgment

This work has been supported by National Institutes of Health grants R35GM147004 and R03OD036491 to G.G.

## Supplemental Materials

### List of Supplemental Materials

- Supplemental Figures
- **Supplemental Figure S1**. Precision and recall of IRR extraction from 540 simulated WGS samples.
- Supplemental Tables
- **Supplemental Table S1**. Regions and motifs of tandem repeat loci used for benchmarking RMSE of ScatTR and state-of-the-art methods
- **Supplemental Table S2**. Regions and motifs of tandem repeat loci with long motifs (21-50 bp) used for benchmarking RMSE of ScatTR and state-of-the-art methods
- **Supplemental Table S2**. Regions and motifs of tandem repeat loci used for benchmarking effect of mutations and sequencing error on ScatTR’s performance

## Supplemental Figures

**Figure S1.**
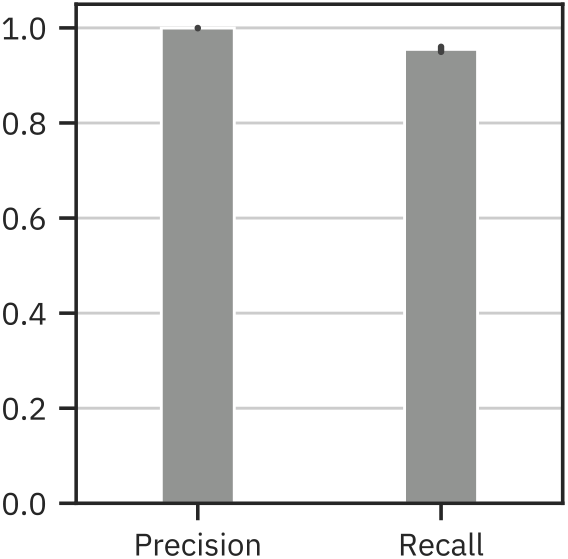
Supplemental Figure S1.Precision and recall of IRR extraction from 540 simulated WGS samples.

## References

Benson, G (Jan. 1999). “Tandem repeats finder: a program to analyze DNA sequences”. eng. In: Nucleic Acids Research 27.2, pp. 573–580. DOI: 10.1093/nar/27.2.573.

Boža, V, B Brejová, and T Vinař (June 2015). “GAML: genome assembly by maximum likelihood”. In: Algorithms for Molecular Biology 10.1, p. 18. DOI: 10.1186/s13015-015-0052-6.

Brook, JD et al. (Feb. 1992). “Molecular basis of myotonic dystrophy: Expansion of a trinucleotide (CTG) repeat at the 3’ end of a transcript encoding a protein kinase family member”. English. In: Cell 68.4. Publisher: Elsevier, pp. 799–808. DOI: 10.1016/0092-8674(92)90154-5.

Budiš, J, M Kucharík, F Ďuriš, J Gazdarica, M Zrubcová, A Ficek, T Szemes, B Brejová, and J Radvanszky (Apr. 2019). “Dante: genotyping of known complex and expanded short tandem repeats”. In: Bioinformatics 35.8, pp. 1310–1317. DOI: 10.1093/bioinformatics/bty791.

Byrska-Bishop, M et al. (Sept. 2022). “High-coverage whole-genome sequencing of the expanded 1000 Genomes Project cohort including 602 trios”. English. In: Cell 185.18. Publisher: Elsevier, 3426–3440.e19. DOI: 10.1016/j.cell.2022.08.004.

Caron, NS, GE Wright, and MR Hayden (1993). “Huntington Disease”. eng. In: GeneReviews®. Ed. by MP Adam, J Feldman, GM Mirzaa, RA Pagon, SE Wallace, LJ Bean, KW Gripp, and A Amemiya. Seattle (WA): University of Washington, Seattle.

Clark, SC, R Egan, PI Frazier, and Z Wang (Feb. 2013). “ALE: a generic assembly likelihood evaluation framework for assessing the accuracy of genome and metagenome assemblies”. In: Bioinformatics 29.4, pp. 435–443. DOI: 10.1093/bioinformatics/bts723.

Cortese, A et al. (Apr. 2019). “Biallelic expansion of an intronic repeat in RFC1 is a common cause of late-onset ataxia”. en. In: Nature Genetics 51.4. Publisher: Nature Publishing Group, pp. 649–658. DOI: 10.1038/s41588-019-0372-4.

Currò, R et al. (May 2024). “Role of the repeat expansion size in predicting age of onset and severity in RFC1 disease”. In: Brain 147.5, pp. 1887–1898. DOI: 10.1093/brain/awad436.

Dashnow, H, M Lek, et al. (Aug. 2018). “STRetch: detecting and discovering pathogenic short tandem repeat expansions”. In: Genome Biology 19.1, p. 121. DOI: 10.1186/s13059-018-1505-2

.Dashnow, H, BS Pedersen, et al. (Dec. 2022). “STRling: a k-mer counting approach that detects short tandem repeat expansions at known and novel loci”. In: Genome Biology 23.1, p. 257. DOI: 10.1186/s13059-022-02826-4.

Dolzhenko, E, MF Bennett, et al. (Dec. 2020). “ExpansionHunter Denovo: a computational method for locating known and novel repeat expansions in short-read sequencing data”. en. In: Genome Biology 21.1, p. 102. DOI: 10.1186/s13059-020-02017-z.

Dolzhenko, E, V Deshpande, et al. (Nov. 2019). “ExpansionHunter: a sequence-graph-based tool to analyze variation in short tandem repeat regions”. In: Bioinformatics 35.22, pp. 4754–4756. DOI: 10.1093/bioinformatics/btz431.

Dolzhenko, E, A English, et al. (May 2023). Resolving the unsolved: Comprehensive assessment of tandem repeats at scale. en. Doi: 10.1101/2023.05.12.540470.

Dolzhenko, E, JJFAv Vugt, et al. (Nov. 2017). “Detection of long repeat expansions from PCR-free whole-genome sequence data”. en. In: Genome Research 27.11, pp. 1895–1903. DOI: 10.1101/gr.225672.117.

Erwin, GS et al. (Jan. 2023). “Recurrent repeat expansions in human cancer genomes”. en. In: Nature 613.7942. Publisher: Nature Publishing Group, pp. 96–102. doi:10.1038/s41586-022-05515-1.

Ghodsi, M, CM Hill, I Astrovskaya, H Lin, DD Sommer, S Koren, and M Pop (Aug. 2013). “De novo likelihood-based measures for comparing genome assemblies”. en. In: BMC Research Notes 6.1, p. 334. DOI: 10.1186/1756-0500-6-334.

Gijselinck, I et al. (Aug. 2016). “The C9orf72 repeat size correlates with onset age of disease, DNA methylation and transcriptional downregulation of the promoter”. en. In: Molecular Psychiatry 21.8. Publisher: Nature Publishing Group, pp. 1112–1124. DOI: 10.1038/mp.2015.159.

Gymrek, M, D Golan, S Rosset, and Y Erlich (June 2012). “lobSTR: A short tandem repeat profiler for personal genomes”. en. In: Genome Research 22.6, pp. 1154–1162. DOI: 10.1101/gr.135780.111.

Gymrek, M, T Willems, et al. (Jan. 2016). “Abundant contribution of short tandem repeats to gene expression variation in humans”. en. In: Nature Genetics 48.1. Number: 1 Publisher: Nature Publishing Group, pp. 22–29. DOI: 10.1038/ng.3461.

Hannan, AJ (May 2018). “Tandem repeats mediating genetic plasticity in health and disease”. en. In: Nature Reviews Genetics 19.5. Publisher: Nature Publishing Group, pp. 286–298. DOI: 10.1038/nrg.2017.115.

Huang, W, L Li, JR Myers, and GT Marth (Feb. 2012). “ART: a next-generation sequencing read simulator”. In: Bioinformatics 28.4, pp. 593–594. DOI: 10.1093/bioinformatics/btr708.

Hunter, JE, E Berry-Kravis, H Hipp, and PK Todd (1993). “FMR1 Disorders”. eng. In: GeneRe-views®. Ed. by MP Adam, J Feldman, GM Mirzaa, RA Pagon, SE Wallace, LJ Bean, KW Gripp, and A Amemiya. Seattle (WA): University of Washington, Seattle.

Kenna, KP et al. (Sept. 2016). “NEK1 variants confer susceptibility to amyotrophic lateral sclerosis”. en. In: Nature Genetics 48.9. Publisher: Nature Publishing Group, pp. 1037–1042. DOI: 10.1038/ng.3626.

Kent, WJ, CW Sugnet, TS Furey, KM Roskin, TH Pringle, AM Zahler, and aD Haussler (June 2002). “The Human Genome Browser at UCSC”. en. In: Genome Research 12.6. Company: Cold Spring Harbor Laboratory Press Distributor: Cold Spring Harbor Laboratory Press Institution: Cold Spring Harbor Laboratory Press Label: Cold Spring Harbor Laboratory Press Publisher: Cold Spring Harbor Lab, pp. 996–1006. DOI: 10.1101/gr.229102.

Kiefer, J (1953). “Sequential minimax search for a maximum”. en. In: Proceedings of the Amer-ican Mathematical Society 4.3, pp. 502–506. DOI: 10.1090/S0002-9939-1953-0055639-3.

Kirkpatrick, S, CD Gelatt, and MP Vecchi (May 1983). “Optimization by Simulated Annealing”. In: Science 220.4598. Publisher: American Association for the Advancement of Science,pp. 671–680. DOI: 10.1126/science.220.4598.671.

Leehey, MA et al. (Apr. 2008). “FMR1 CGG repeat length predicts motor dysfunction in pre-mutation carriers”. In: Neurology 70.16 part 2. Publisher: Wolters Kluwer, pp. 1397–1402. DOI: 10.1212/01.wnl.0000281692.98200.f5.

Logsdon, GA et al. (Sept. 2024). Complex genetic variation in nearly complete human genomes. en. Pages: 2024.09.24.614721 Section: New Results. DOI: 10.1101/2024.09.24.614721.

Martorell, L, MA Pujana, V Volpini, A Sanchez, J Joven, E Vilella, and X Estivill (1997). “The repeat expansion detection method in the analysis of diseases with CAG/CTG repeat expansion: Usefulness and limitations”. en. In: Human Mutation 10.6. eprint: https://onlinelibrary.wiley.com/doi/1004%281997%2910%3A6%3C486%3A%3AAID-HUMU11%3E3.0.CO%3B2-W, pp. 486–488. DOI: 10.1002/(SICI)1098-1004(1997)10:6<486::AID-HUMU11>3.0.CO;2-W.

Mousavi, N, S Shleizer-Burko, R Yanicky, and M Gymrek (Sept. 2019). “Profiling the genome-wide landscape of tandem repeat expansions”. In: Nucleic Acids Research 47.15, e90. DOI: 10.1093/nar/gkz501.

Nishimura, D (Apr. 2000). “RepeatMasker”. In: Biotech Software & Internet Report 1.1-2. Publisher: Mary Ann Liebert, Inc., publishers, pp. 36–39. DOI: 10.1089/152791600319259.

Nurk, S et al. (Apr. 2022). “The complete sequence of a human genome”. In: Science (New York, N.Y.) 376.6588, pp. 44–53. DOI: 10.1126/science.abj6987.

Rahman, A and L Pachter (Jan. 2013). “CGAL: computing genome assembly likelihoods”. en.In: Genome Biology 14.1, R8. DOI: 10.1186/gb-2013-14-1-r8.

Siddique, N and T Siddique (1993). “Amyotrophic Lateral Sclerosis Overview”. eng. In: GeneRe-views®. Ed. by MP Adam, J Feldman, GM Mirzaa, RA Pagon, SE Wallace, LJ Bean, KW Gripp, and A Amemiya. Seattle (WA): University of Washington, Seattle.

Song, JHT, CB Lowe, and DM Kingsley (Sept. 2018). “Characterization of a Human-Specific Tandem Repeat Associated with Bipolar Disorder and Schizophrenia”. In: The American Journal of Human Genetics 103.3, pp. 421–430. DOI: 10.1016/j.ajhg.2018.07.011.

Tang, H et al. (Nov. 2017). “Profiling of Short-Tandem-Repeat Disease Alleles in 12,632 Human Whole Genomes”. In: The American Journal of Human Genetics 101.5, pp. 700–715. doi:10.1016/j.ajhg.2017.09.013.

Tankard, RM, MF Bennett, P Degorski, MB Delatycki, PJ Lockhart, and M Bahlo (Dec. 2018). “Detecting Expansions of Tandem Repeats in Cohorts Sequenced with Short-Read Sequencing Data”. In: The American Journal of Human Genetics 103.6, pp. 858–873. doi:10.1016/j.ajhg.2018.10.015

Tassone, F, J Adams, EM Berry-Kravis, SS Cohen, A Brusco, MA Leehey, L Li, RJ Hagerman, and PJ Hagerman (June 2007). “CGG repeat length correlates with age of onset of motor signs of the fragile X-associated tremor/ataxia syndrome (FXTAS)”. eng. In: American Journal of Medical Genetics. Part B, Neuropsychiatric Genetics: The Official Publication of the International Society of Psychiatric Genetics 144B.4, pp. 566–569. DOI: 10.1002/ajmg.b.30482.

Treangen, TJ and SL Salzberg (Jan. 2012). “Repetitive DNA and next-generation sequencing: computational challenges and solutions”. en. In: Nature Reviews Genetics 13.1. Publisher: Nature Publishing Group, pp. 36–46. DOI: 10.1038/nrg3117.

Trost, B et al. (Oct. 2020). “Genome-wide detection of tandem DNA repeats expanded in autism”. In: Nature 586.7827, pp. 80–86. DOI: 10.1038/s41586-020-2579-z.

Willems, T, D Zielinski, J Yuan, A Gordon, M Gymrek, and Y Erlich (June 2017). “Genome-wide profiling of heritable and de novo STR variations”. en. In: Nature Methods 14.6, pp. 590–592. DOI: 10.1038/nmeth.4267.

Zhou, ZD, J Jankovic, T Ashizawa, and EK Tan (Mar. 2022). “Neurodegenerative diseases associated with non-coding CGG tandem repeat expansions”. en. In: Nature Reviews Neurology 18.3. Publisher: Nature Publishing Group, pp. 145–157. doi:10.1038/s41582-021-00612-7.

